# High Variability Periods in the EEG: A New Temporal Metric that Reflects Brain States

**DOI:** 10.1101/2022.06.27.497770

**Authors:** Dhanya Parameshwaran, Tara Thiagarajan

## Abstract

Here we define a new metric to characterize temporal patterns of amplitude variability in the EEG signal and demonstrate that this feature of the signal varies significantly with brain state. The metric uses the standard deviation of waveform amplitude in a short moving time window of 3-20 seconds with 50% overlap. We define “High Variability Periods” or “HVPs” as segments when the moving standard deviation of the waveform amplitude is continuously higher than a cutoff defined as the 25th percentile of variability. HVPs typically last on the order of tens of seconds and are punctuated by low variability periods or “LVPs” of shorter duration. We further show that the HVP characteristics differ between various conditions. HVPs in the resting state eyes closed condition are substantially and significantly shorter in duration and smaller in area relative to eyes open. In addition, in recordings from monkeys, HVPs disappear under anesthesia and do not reappear in early periods of recovery from anesthesia suggesting long term changes to the signal. Altogether this demonstrates that HVP metrics have discriminatory power and may therefore be important in predicting brain states and outcomes. Finally, they are fast to estimate and can be tracked in time, and therefore suited for near real-time monitoring in low-electrode configuration systems.

## Introduction

The primary advantage of EEG relative to other neuroimaging methods is its high temporal resolution. However, there are relatively few metrics that exploit or effectively describe the temporal properties of the signal. Commonly used metrics utilize computations and transformations that either ignore temporal aspects of the signal (e.g. spectral metrics) or apply numerous transformations that are based on assumptions of time scales or aspects of underlying behavior of the brain (e.g. entropy metrics). Thus there is a need for new approaches to the EEG signal that provide insights into the temporal structure of the EEG signal.

In addition, few metrics lend themselves easily to real-time monitoring by either requiring high density electrode configurations, significant computational power, or simply not being structured for high resolution temporal tracking. Thus expanding the application of EEG requires metrics that are computationally fast, easy to implement with low electrode configurations and can provide a meaningful view of changes in time.

Finally, while there are numerous behavioral conditions and brain states, individual metrics have few degrees of freedom resulting in a one-to-many mapping of the EEG characteristic to outcomes. Thus multiple orthogonal perspectives of the signal are needed to provide deeper insights into the relationship between brain states and outcomes.

Here we propose a new metric that characterizes High Variability Periods (HVPs) in the EEG and meets many of the conditions above. This metric provides a view of the temporal variability of the amplitude of the signal, is computationally fast, and can be applied on a single channel basis. Furthermore, it makes no assumptions about the underlying behavior of the brain and is uncorrelated with existing entropy and complexity metrics that are commonly used to estimate aspects of variability in the signal. Finally, we show that this metric varies by condition and mental state using various examples from humans and monkeys indicating its utility as a biomarker of brain states.

## Methods

### Human EEG Recordings

Our demonstration of the computation and properties of this metric utilize EEG recordings obtained from 50 adults between the ages of 21 and 50 as previously described in [1]. EEG recordings were obtained while participants were sitting quietly for three minutes with their eyes closed (EC) or with their eyes open and looking at images on a laptop screen (EO). Briefly, the recordings were obtained using the Emotiv EPOC+ which utilizes a bilateral mastoid reference (M1, a ground reference point for measuring the voltage of the other sensors and M2, a feed-forward reference point for reducing electrical interference from external sources). The EPOC+ signals were high-pass filtered with a 0.16 Hz cut-off, pre-amplified and low-pass filtered at an 83 Hz cut-off. The analogue signals were then digitized at 2048 Hz and filtered using a 5th-order sinc notch filter (50 and 60 Hz) before being down-sampled to 128 Hz (company communication). The effective bandwidth of the signal is therefore between 0.16 and 45 Hz. All recordings were obtained following informed written consent using protocols approved by the Health Media Lab’s Institutional Review Board (IRB #00001211, IORG #0000850) in accordance with the requirements of the US Code of Federal Regulations for the Protection of Human Subjects and Sigma IRB (India).

### Monkey EEG Recordings

EEG and ECOG simultaneous recordings from monkeys were obtained from Neurotycho, an open-access collection of multidimensional, invasive EEG recordings from multiple macaque monkeys [2, 3]. The EEG signal was recorded in two monkeys from 19 channels with a sampling rate of 4096 Hz. The location of the EEG electrodes was determined by a 10-20 system without Cz using the NeuroPRAX system. EEG signals were referenced to an average between the signals recorded from the bilateral mastoids. Recordings were down-sampled to 128 Hz to match the human recordings.

For the anesthesia experiment, EEGs were recorded after injection of an anesthetic agent and an antagonist. The monkey was seated in a primate chair with the movements of both arms and the head restricted and heart rate and respiration were monitored during the entire experiment. The anesthetic agent was a combination of ketamine (8.21 mg/kg for M1 and 5.00 mg/kg for M2) and medetomidine (0.05 mg/kg for M1 and 0.02 mg/kg for M2). Data was collected in 5-minute blocks. In the low-anesthetic state, 5 min of EEG was recorded shortly after the injection. In the deep-anesthetic state, 5 min of EEG data was recorded when sneeze, grasp, and blink reflexes had vanished. In the recovery state, 5 min of EEG was recorded shortly after injection of the antagonist (atipamezole, 0.21 mg/kg for M1 and 0.09 mg/kg for M2).

### Defining High Variability Periods (HVPs)

HVPs are identified by first computing the standard deviation of the amplitude values (A_SD) for each segment or window of the signal of length *m*, moving the window along the signal duration with a 50% overlap. Figure 1A and C shows the EEG signal (grey) normalized to the overall channel A_SD as well as the A_SD values for three different values of m (3, 9 and 15 seconds) shifted with a 50% overlap for one human and one monkey respectively. The amplitude value corresponding to the 25th percentile value of the A_SD values computed across all windows across the full length of the signal is then used as a threshold T (dotted line in Figure 1B, D). High Variability Periods or HVPs are then defined as those stretches where the A_SD values are continuously above the threshold T. Conversely Low Variability Periods or LVPs are defined as those stretches where the A_SD values are continuously *below* the threshold T. While Figures 1A and 4A,B show the EEG signal normalized by A_SD value in order to accommodate both the signal and moving window on the same graph, other figures utilize the amplitude SD value in μV corresponding to the SD threshold.

**Figure 1:**
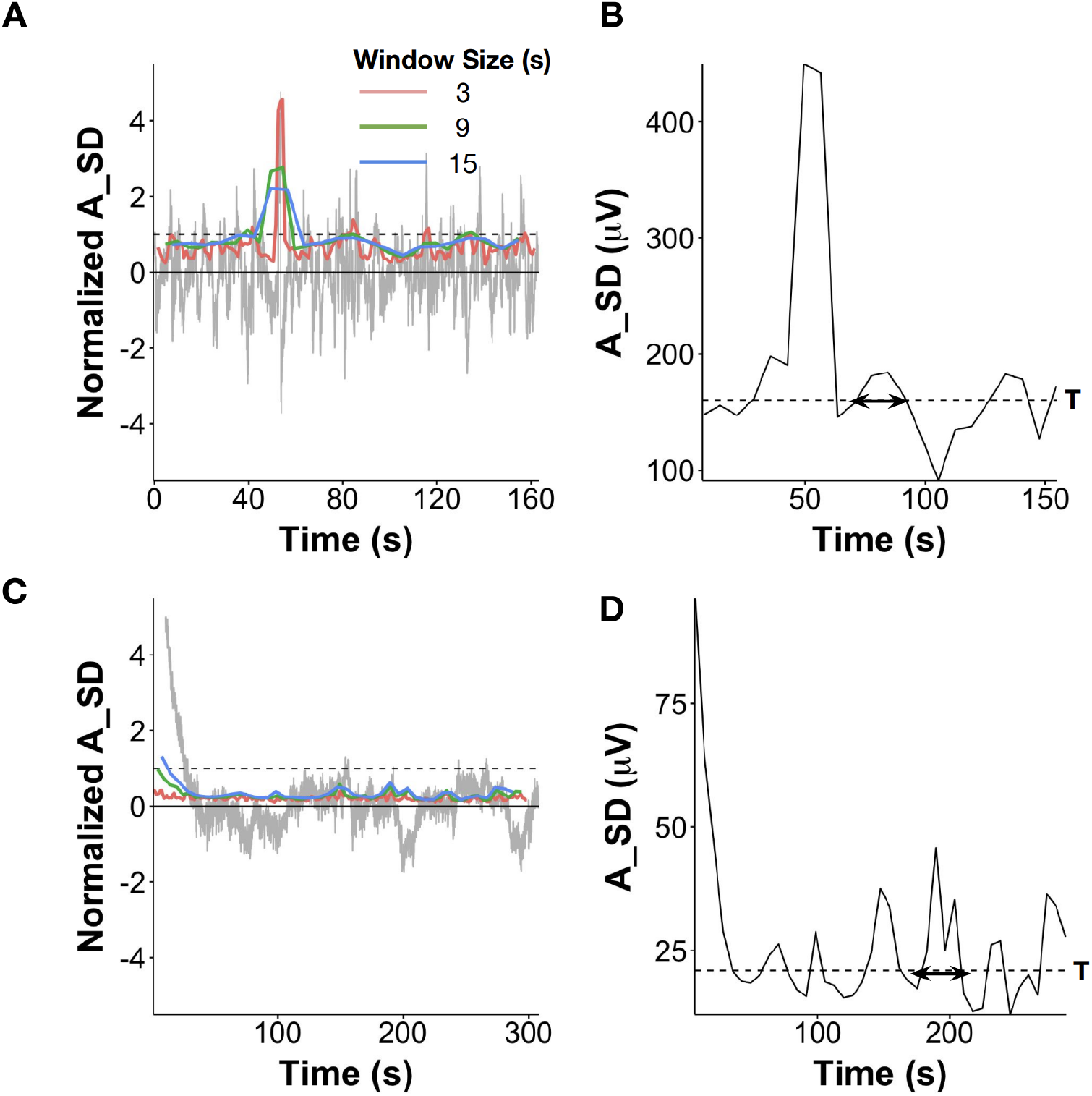
Definition of High Variability Periods (HVP) A. Resting state EEG signal from one person with amplitude shown in units of standard deviation (A_SD, grey) along with the SD of the signal amplitude estimated for varying window sizes from 3, 9 and 15 to 30 seconds (colored lines). Dashed line represents the SD of the signal amplitude of the whole trace i.e. A_SD or 1 SD of the amplitude. B. The moving A_SD with a 15 second window and 50% overlap for the EEG signal as shown in A. The dashed line represents the 25th percentile of the moving A_SD value which serves as the threshold T. All segments >T are defined as High Variability Periods or HVPs. Segments consistently <T are defined as low variability periods or LVPs. Durations are computed as the time between the threshold crossings. Area (for HVPs only) is computed as the sum of all SD values across the duration D. C. Similar resting state signal along with SD estimates for different moving window sizes as shown in A, but from a monkey rather than human. D. The moving A_SD with a 15 second window and 50% overlap for the EEG signal as shown in C.

### Computing High Variability Periods or HVP Metrics

We compute three metrics that describe these High Variability Periods. 1) The average and/or median HVP duration (HVP-D) computed as the average or median width of all HVPs across the recording, 2) The average and/or median HVP area (HVP-A), computed as the average or median of the sum of all values across the HVP period and 3) The HVP rate (HVP-R) which is the number of HVPs per minute of EEG recording. Conversely Low Variability Periods (LVPs) are defined as those segments continuously less than the threshold T for which durations were also computed.

We note that when comparing multiple conditions within the same recording (i.e. resting Eyes Open and Eyes Closed), the threshold was determined using the resting Eyes Closed (EC) in all conditions (Figures 2 and 3). When comparing multiple states in anesthesia (Figure 4), we used a 3s window size to compute the A_SD. Furthermore, the 25th percentile threshold T used to compute HVPs was determined based on the initial rest condition and applied to all subsequent states of low and deep anesthesia and recovery. Mean values were computed by first averaging across channels for each individual within a group and then computing the mean and standard error of these averages across all individuals.

**Figure 2:**
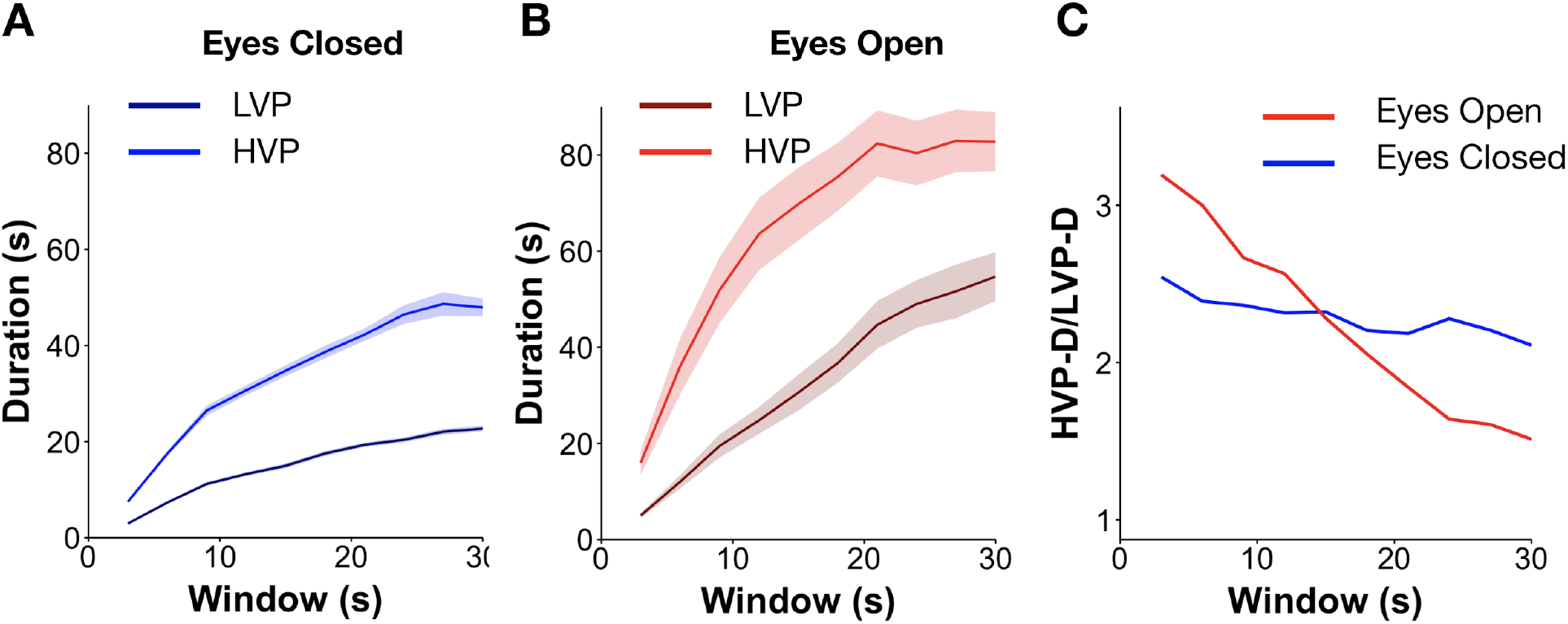
Impact of window size on High Variability Periods (HVPs) A. Mean duration of HVPs and LVPs across all human participants (mean ±SEM) as a function of window size in the eyes closed state (EC). B. Mean duration of HVPs and LVPs as in A in the eyes open rest state (EO). C. Ratio of HVP-duration to LVP-duration across window sizes in EO and EC conditions.

**Figure 3:**
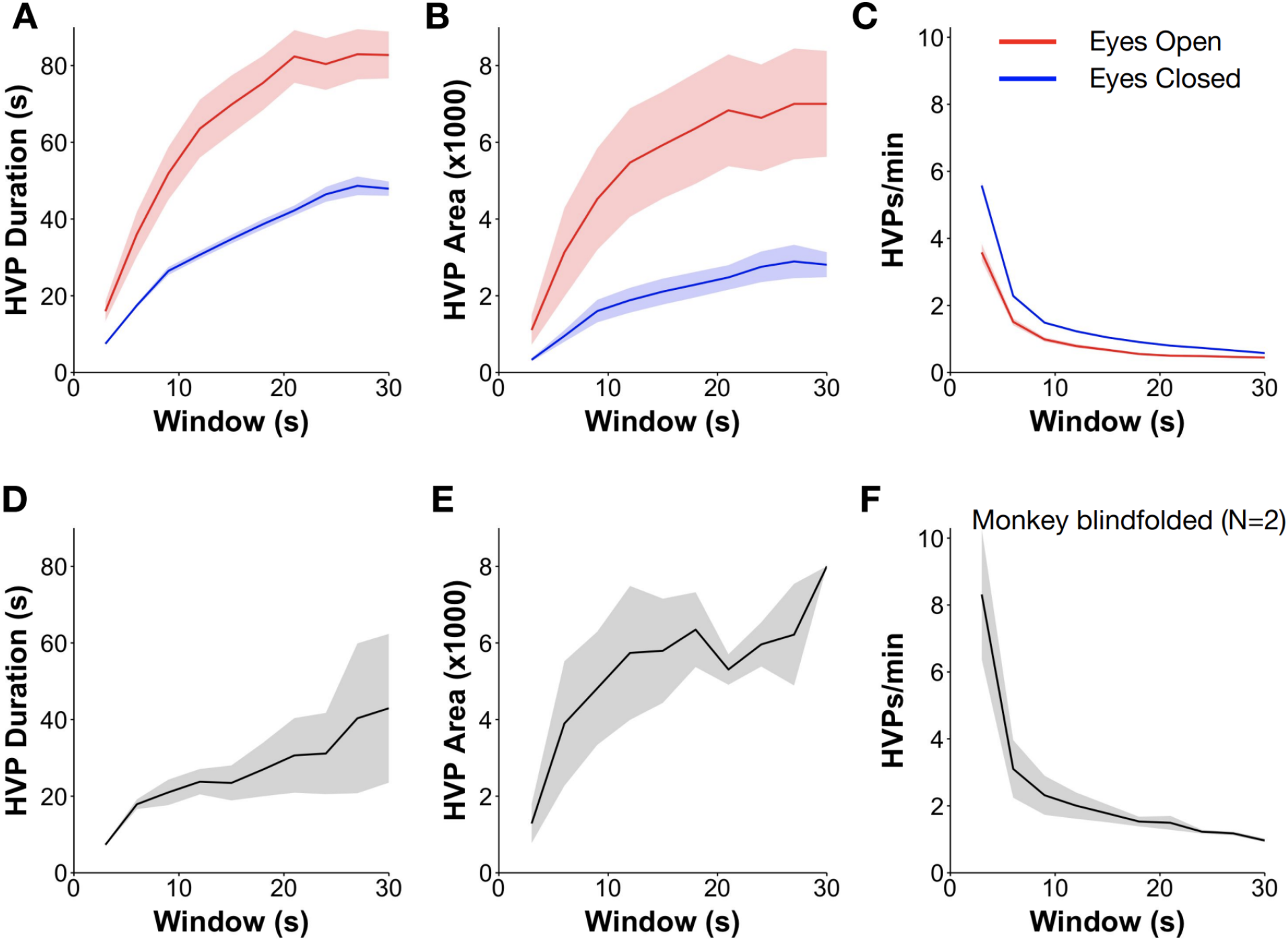
HVP metrics across conditions A. HVP duration across all human participants (mean ±SEM) for eyes closed (EC; blues) and eyes open conditions (EO; reds) as a function of window size. B. HVP area across all participants (mean ±SEM) for the conditions as in A. C. HVP rate across all participants (mean ±SEM) for the conditions as in A. D. HVP duration as in A averaged across both monkeys (blindfolded). E. HVP area averaged across both monkeys as in B F. HVP rate averaged across both monkeys as in C

**Figure 4:**
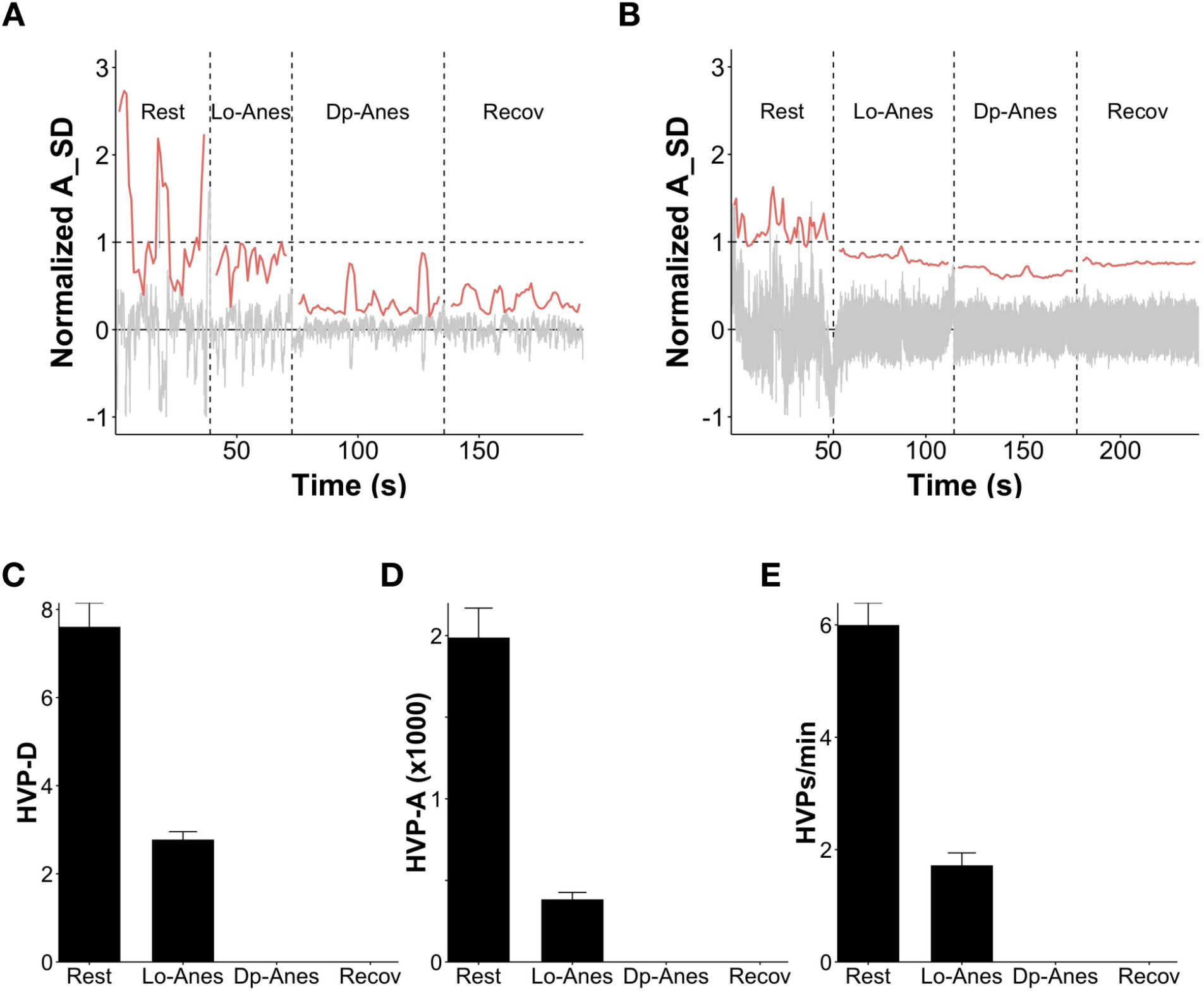
Impact of anesthesia on HVPs in the EEG signal A-B. Examples from a single channel in each of two monkeys showing the EEG signal (gray, normalized by the SD) before (rest) and after injection of ketamine (low-anesthetic and deep-anesthetic) and following injection of an antagonist for reversal of the anesthetic effect (recovery). The moving A_SD used to calculate HVPs shown in red. The recording in Figure A used a resting state threshold of 280 μV and the recording in Figure B used a resting state threshold of 153 μV. C-E. HVP duration, area and rate per minute in each state of anesthesia. HVP events completely disappeared in deep anesthesia, and did not reappear in the initial recovery phase.

A function for computing HVP is available on request.

### Computing of Entropy and Waveform Complexity Metrics

Sample Entropy, Lempel Ziv Entropy and Waveform Complexity were computed as described in [4].

## Results

### Window size and HVP characteristics

We first looked at how the durations of HVPs and LVPs varied as a function of window size in the eyes open and eyes closed conditions in humans (Figure 2; Humans N=50) and in monkeys blindfolded (Figure 3D, N=2). Across both species, and conditions HVP durations (HVP-D) were on the order of tens of seconds and increased sub-linearly with the window size. The mean value of the 25th percentile A_SD of each state was used as a threshold to estimate HVP and LVP metrics. In the eyes closed condition the ratio of HVPs to LVPs ranged from 2.1 to 2.5 across all window sizes (Figure 2A). However, in the eyes open condition (EO), HVPs were on average 2.4 times and up to 3.2 times longer in duration than LVPs across the range of window sizes (Table 1, Figure 2B).

**Table 1:** HVP and LVP metrics across Eyes Open and Eyes Closed conditions

| EyesClosed |  |  |  |  |  |  |  |
| --- | --- | --- | --- | --- | --- | --- | --- |
| m | HVP-D | HVP-A | HVP-R | LVP-D | LVP-A | LVP-R | HVP-D/LVP-D |
| 3 | 7 | 332 | 5.6 | 3 | 51 | 3.4 | 2.5 |
| 6 | 18 | 953 | 2.3 | 7 | 167 | 1.2 | 2.4 |
| 9 | 26 | 1600 | 1.5 | 11 | 300 | 0.7 | 2.4 |
| 15 | 35 | 2109 | 1.0 | 15 | 455 | 0.6 | 2.3 |
| 21 | 42 | 2478 | 0.8 | 19 | 607 | 0.4 | 2.2 |
| 30 | 48 | 2810 | 0.6 | 23 | 752 | 0.4 | 2.1 |

*Table 1: HVP and LVP metrics across Eyes Open and Eyes Closed conditions*
| EyesOpen |  |  |  |  |  |  |  |
| --- | --- | --- | --- | --- | --- | --- | --- |
| m | HVP-D | HVP-A | HVP-R | LVP-D | LVP-A | LVP-R | HVP-D/LVP-D |
| 3 | 16 | 1107 | 3.6 | 5 | 90 | 2.8 | 3.2 |
| 6 | 36 | 3133 | 1.5 | 12 | 287 | 1.2 | 3.0 |
| 9 | 52 | 4519 | 1.0 | 19 | 512 | 0.8 | 2.7 |
| 15 | 70 | 5929 | 0.7 | 31 | 884 | 0.6 | 2.3 |
| 21 | 82 | 6834 | 0.5 | 45 | 1268 | 0.5 | 1.8 |
| 30 | 83 | 7001 | 0.4 | 55 | 1572 | 0.4 | 1.5 |

### Differences in HVP characteristics between groups

HVPs were significantly higher in both duration (HVP-D) and area (HVP-A) in the eyes open condition relative to the eyes closed condition for all window sizes (Table 1, Figures 3A, B), and increased sub-linearly with window size in all conditions and in both monkeys and humans (Figures 3B,E). Conversely, the rates of HVPs (HVP-R) which were generally less than 6 HVPs/minute decreased exponentially with window size (Figure 3C,F). Overall, HVP-D in eyes open was about twice as long as in the eyes closed condition for all window sizes while HVP-A was on average three times as long in the eyes open condition (Table 2).

**Table 2:**
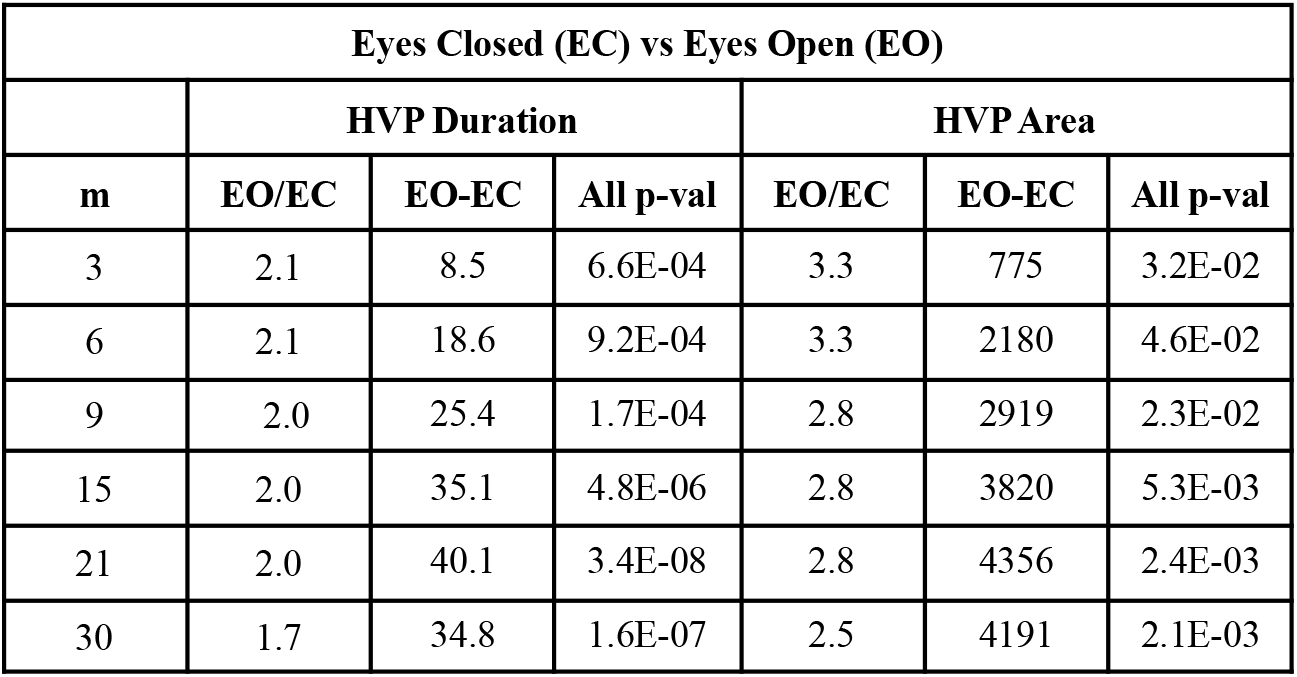
Differences in HVP metrics between EC and EO conditions

| Eyes Closed (EC) vs Eyes Open (EO) |  |  |  |  |  |  |
| --- | --- | --- | --- | --- | --- | --- |
|  | HVP Duration |  |  | HVP Area |  |  |
| m | EO/EC | EO-EC | All p-val | EO/EC | EO-EC | All p-val |
| 3 | 2.1 | 8.5 | 6.6E-04 | 3.3 | 775 | 3.2E-02 |
| 6 | 2.1 | 18.6 | 9.2E-04 | 3.3 | 2180 | 4.6E-02 |
| 9 | 2.0 | 25.4 | 1.7E-04 | 2.8 | 2919 | 2.3E-02 |
| 15 | 2.0 | 35.1 | 4.8E-06 | 2.8 | 3820 | 5.3E-03 |
| 21 | 2.0 | 40.1 | 3.4E-08 | 2.8 | 4356 | 2.4E-03 |
| 30 | 1.7 | 34.8 | 1.6E-07 | 2.5 | 4191 | 2.1E-03 |

The HVP duration and area in monkeys followed a similar pattern. Overall HVP durations were slightly lower or similar at all window sizes for these blindfolded monkeys compared to the eyes closed state in humans (Figure 3D-E). While this suggests the possibility of interesting inter-species differences, since the number of monkeys were only 2 and recording devices are different, it is not possible to make a statistical or reliable comparison between monkeys and humans.

For an inter-species comparison, more directly comparable methodologies would need to be employed with a larger number of monkeys to draw reliable conclusions.

### Effect of anesthesia on HVP metrics

We next looked at how HVPs behaved under ketamine anesthesia in each of two monkeys (Figure 4) in recordings of 5 minute blocks before anesthesia (rest), immediately after (two successive blocks called low and deep anesthesia) and after injection of a reversing antagonist (recovery). Here we show that HVPs reduced from 6/minute during the rest block to 1.7 HVPs/minute during the low-anesthetic block and then completely disappeared in deep anesthesia, and did not reappear in the initial recovery phase. Those HVPs that persisted in low anesthesia were more than three times smaller in duration and area (Figure 3C-E).

### Relationship to other temporal metrics

Other temporal metrics such as entropy and complexity also quantify variability in the signal. While HVP metrics are different in that they provide a view of variability in time rather than a view of the overall variability of the signal, the novel utility of HVPs depends on the metric providing substantially different information about the signal relative to these other temporal metrics. We therefore looked at the relationship between various entropy and complexity metrics, showing that Sample Entropy, Liv Zempl Complexity and Waveform Complexity are at best very weakly and not significantly correlated with the HVP metrics of duration, area and rate for the same signal (N=28, Table 3). Thus HVP metrics provide a substantially distinct view of the signal relative to these other metrics with novel utility.

**Table 3:** Correlations between HVP metrics and Entropy/Complexity measures

| HVP metrics | Entropy/Complexity metrics | Pearson's r | p-val |
| --- | --- | --- | --- |
| Duration | Lempel-Ziv Complexity | 0.01 | 0.97 |
| Duration | Waveform Complexity | 0.17 | 0.42 |
| Duration | Sample Entropy | -0.09 | 0.68 |
| Area | Lempel-Ziv Complexity | 0.06 | 0.76 |
| Area | Waveform Complexity | -0.18 | 0.40 |
| Area | Sample Entropy | -0.32 | 0.12 |
| Rate | Lempel-Ziv Complexity | 0.01 | 0.96 |
| Rate | Waveform Complexity | -0.18 | 0.38 |
| Rate | Sample Entropy | 0.10 | 0.65 |

## Discussion

### Characteristics of the HVP metric

Here we describe a new metric that characterizes periods of high amplitude variability in the signal. This metric is computationally light and is computed at the level of a single channel to provide a view of the temporal variability. It takes advantage of the high temporal resolution of the signal and does not make any assumptions about the underlying source or otherwise transform the signal based on any assumptions about the underlying behavior of the brain. Rather it is simply a readout of the temporal amplitude variability. The most important parameter is the threshold of A_SD used for definition of the HVP. For consistency we suggest maintaining the 25th percentile as the threshold rather than varying this parameter in order to prevent inclusion of small changes that may arise from noise. The segment of signal is used to compute the A_SD value and therefore the threshold used to define HVPs is of paramount consideration. Here we have pegged everything to the eyes-closed rest state using this threshold for eye-open or anesthesia conditions in order to make an appropriate comparison of the two states. Note that if the threshold was computed separately for the eyes open condition, the results for HVP-D and HVP-A would be different since the A_SD value would be much higher than in the eyes-closed condition and therefore the threshold would be higher as well.

The only other significant parameter choices are the window size and overlap and any use of this metric should therefore clearly specify window both. While varying the window size may be beneficial when working with signals of different durations, maintaining the overlap at 50% is advised.

### Utility of the metric

We have demonstrated that this characteristic of the signal (HVPs) differs significantly between the conditions of eyes open and eyes closed and is suppressed by anesthesia. Thus we suggest that it might have broad application in monitoring brain states under various treatments and therapies, and could provide insight into the nature of mental activity being performed.

Its suppression by anesthesia but lack of recovery immediately after removal of the anesthetic suggest that it might reflect cognitive state rather than consciousness per se and might be investigated in the context of conditions such as postoperative cognitive impairment which has been commonly described [5].

Finally, the additional utility of the HVP metric rests largely on whether it can provide an additional view of the signal not captured by other metrics. In particular, various entropy and complexity measures also indirectly characterize temporal variability in the signal, and like HVPs, Sample Entropy has previously been shown to decrease with anesthesia [6, 7]. Here we have shown that all HVP metrics are largely uncorrelated to these entropy and complexity measures. Thus they provide a novel perspective that can serve as a biomarker of brain states as well as expand predictive capability of brain states when used in conjunction with other metrics.

